# Metal-induced abnormalities in diatom girdle bands

**DOI:** 10.1101/501619

**Authors:** Adriana Olenici, Saúl Blanco, María Borrego-Ramos, Francisco JiméNez-GóMez, Francisco Guerrero, Laura Momeu, Călin Baciu

## Abstract

There have been a number of studies that described a serial of type of teratology occurring in different diatom taxa and that highlight the relation between metal concentration and diatom deformities, but this subject still remain not deeply understood. The present study refers to the effect of metal pollution on the diatom *Achnanthidium minutissimum* s.l. by describing a new form of teratology. The samples were collected in a mine area, Rosia Montana, from Romania. We observed that, exposed to environmental stress, the frustule of diatom cells appeared altered in several ways, with abnormal forms occurring in different diatom species that presented deformed valve outlines, modifications of the raphe canal system, irregular striation or mixed teratologies. In a particular sampling location where *A. minutissimum* s.l. was identified as the dominant species, 20.53% of the individuals presented an unreported type of deformity. This kind of teratology affects the cingulum, the valvocopula more exactly, by becoming markedly undulate.

## 1. Introduction

Diatoms are unicellular microorganisms with a widespread distribution in world aquatic and terrestrial ecosystems. Their most distinctive characteristic is the presence of a siliceous cell wall called the frustule, composed of two units (the valves) joint by a number of connective bands that constitute the cingulum. This cingulum allows the expansion of the frustule to accommodate the formation of new valves and bands during cell division. During asexual reproduction, new girdle bands are deposited on the cell surface through exocytosis to allow cell expansion (De Sanctis et al., 2016). Secretion of the first girdle band indicates the beginning of a new cell cycle during which the cell grows by increasing the distance between the valves (Kröger and Wetherbee, 2000). Silica deposition to form new girdle bands occurs within membrane-bound vesicles (silica deposition vesicles or SDVs) within the protoplast, microfilaments (actin fibers) being also closely associated with forming these structures (Cox, 2011). Each girdle band is produced in an individual SDV (Kröger and Poulsen, 2007). Depending on the species, girdle band formation can occur in different phases during cell cycle, either before or after cytokinesis (Lechner and Becker, 2015). Contrary to valve features, this cingulum has largely been ignored in studies of diatom taxonomy and morphology (Johnson and Rosowski, 1992).

Apart from natural morphological variation, it has been largely recognized the occurrence of teratological forms in diatom populations, i.e. abnormal cells that differ in shape and/or frustular features. Teratological forms have been defined as non-adaptive phenotypic abnormalities usually involving the outline of valves or their striation pattern (Falasco et al., 2009b). Mechanisms inducing teratologies are not fully understood, although they may express short-term phenotypic responses, problems with gene expression (i.e., assembly line malfunction) or true alterations in the genes (Lavoie et al., 2017). According to Cattaneo et al. (2004) and Smol (2008), diatom morphological alterations could be considered a tool for monitoring environmental impairment, and there have been a number of studies that highlight the relation between metal concentration and diatom teratology and that described a serial of type of deformities occurring in different taxa (Falasco et al., 2009a, b; Morin et al., 2008a, b; Lavoie et al. 2012). In any case, the impact of pollution on diatom morphology should be regarded as a combined response to the presence of different stressors in the environment (Falasco et al., 2009a), and seems to depend on the genus involved (Martin-Jézéquel and Lopez, 2003).

## 2. Materials and Methods

Within the types of deformations acknowledged in the literature (chiefly: irregular valve outline, atypical sternum/raphe, and aberrant stria/areolae patterns, see Lavoie et al., 2017), abnormalities affecting girdle bands have not been described to date, although Falasco et al. (2009a) mentioned alterations occurring on the girdle bands in *Staurosira venter* and *Aulacoseira italica*. In this context, this note accounts for the effect of metal pollution on the diatom *Achnanthidium minutissimum* s.l. by describing this new form of teratology. The samples were collected in Abrud River (N46°15’/E23°5’) in a mine area named Rosia Montana (Romania) and were processed following European standards (EN 14407, 2004). The biologic material was collected from the surface of submerged stones in the flow, in the euphotic zone of the river, using a toothbrush, and the sample was preserved in 4% v/v formaldehyde. By oxidizing organic matter with hot hydrogen peroxide 30% v/v clean frustules have been obtained, and permanent microscopic slides were mounted using a refractive resin (colophony - Sunchemy International Co. Ltd, RI ∼ 1.7). At least 400 valves were identified and counted using an Olympus BX 60 microscope, according to usual taxonomic references (Hofmann et al., 2011 and references therein).

## 3. Results and Discussion

We observed that, exposed to environmental stress, the frustule of diatom cells appeared altered in many ways, with abnormal forms occurring in different diatom species that presented deformed valve outlines, modifications of the raphe canal system (displaced fibulae), irregular striation or mixed teratologies (different types of deformed valves and abnormal central area locations or irregular striation). In a particular sampling location (N46°15’39,90”/E23°5’11,80 “), where *A. minutissimum* s.l. was identified as the dominant species, 20.53% of the individuals presented an unreported type of deformity. This kind of teratology affects the cingulum, the valvocopula more exactly (the first of the girdle bands, attached to the valve), by becoming markedly undulate (as shown in Figure 1, 2 and 3). The deformity was observed using both optical and scanning electron microscopy, in processed and unprocessed samples. In comparison with the other sampling sites from the study area, heavy metal and ionic concentrations were lower in this site, although NO_3^-^_, SO_4^2-^_ and Pb levels indicate a moderate to bad water quality according to Romanian standards (Ord. 161, 2006). We have recently demonstrated (Olenici et al., 2017) that the occurrence of teratological diatoms in this area affected by acid mine drainage can be directly linked to the presence of heavy metals; particularly Zn concentrations correlated positively with the degree of deformation in *Achnanthidium* valve outline. Zinc is known to produce asymmetrical, abnormal, and bent frustules (Lavoie et al., 2017), specifically in *A. minutissimum* (Cantonati et al., 2014).

**Figure 1.**
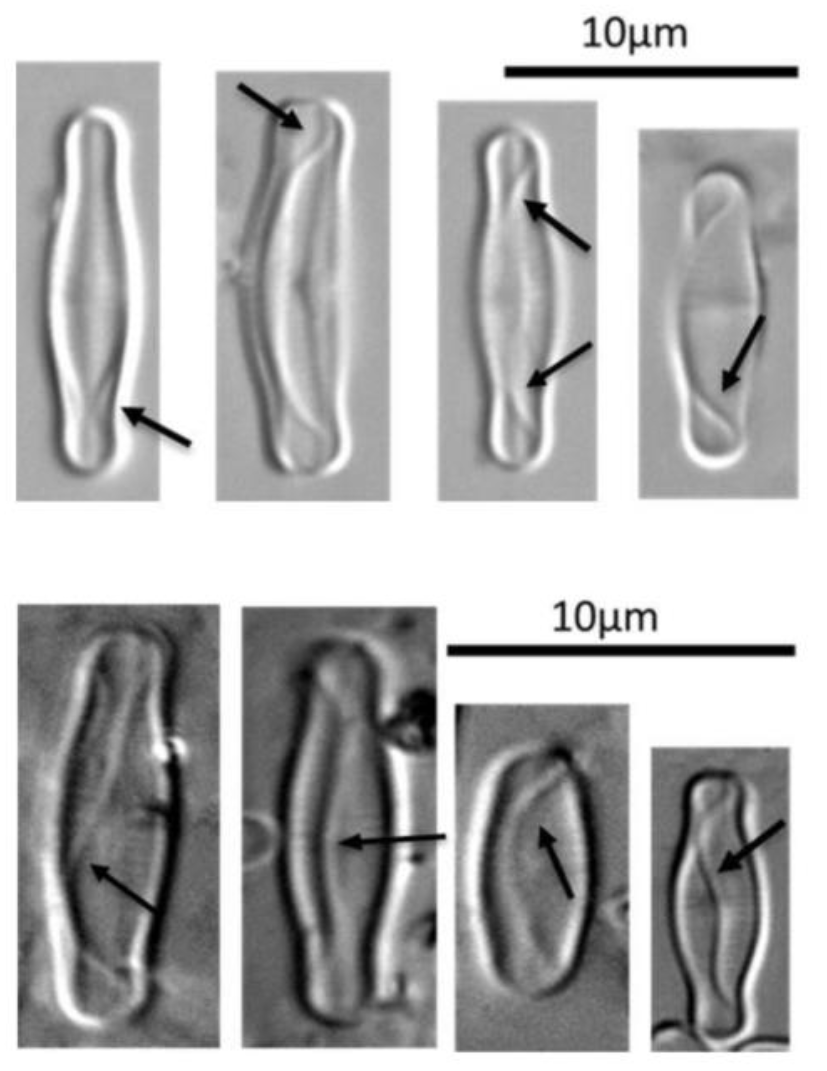
Deformed valvocopula seen at the optical microscope.

**Figure 2.**
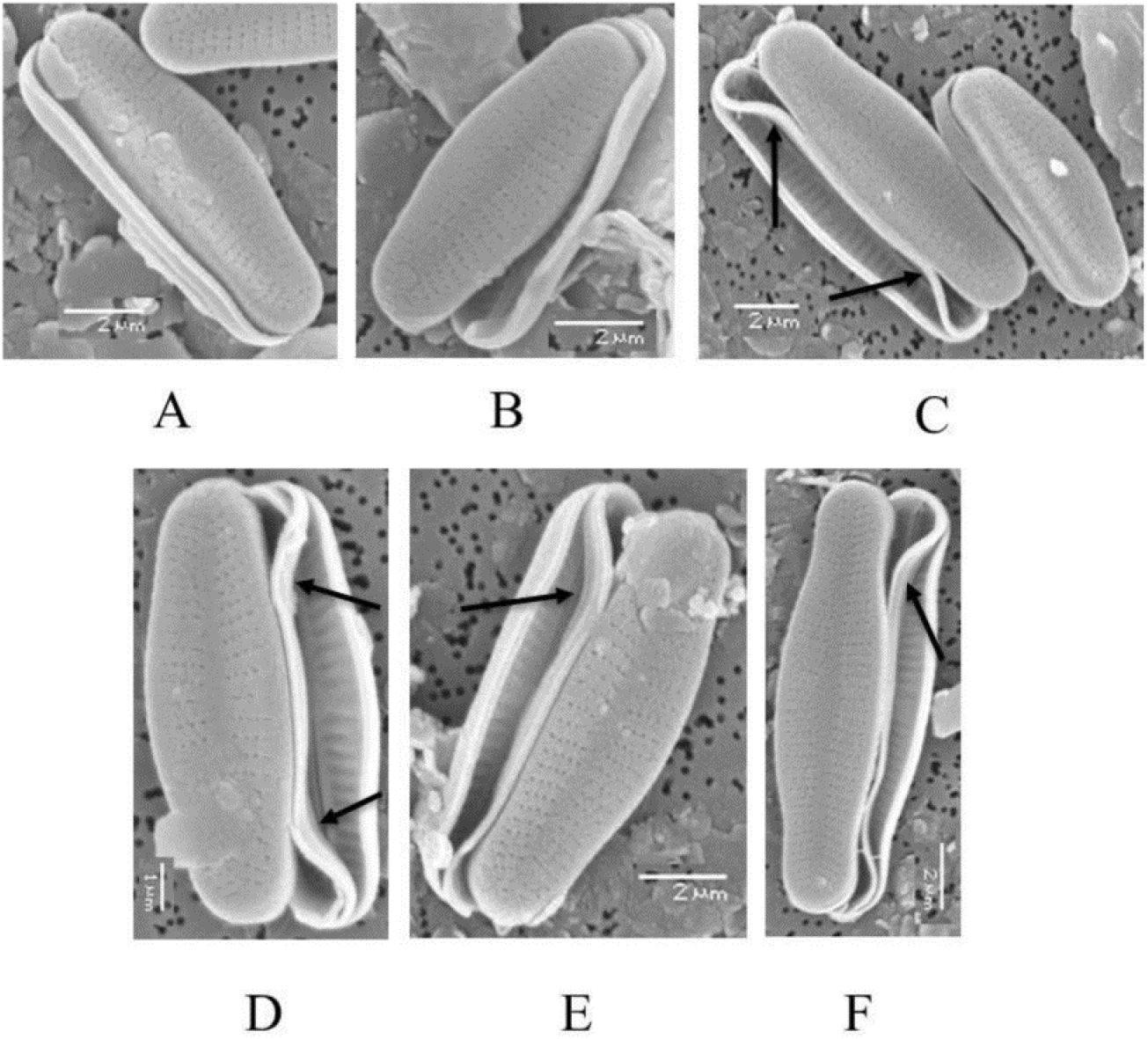
Deformed valvocopula seen at SEM by comparing with normal one (A and B = normal frustules; C, D, E and F = abnormal frustules identified in processed sample).

**Figure 3.**
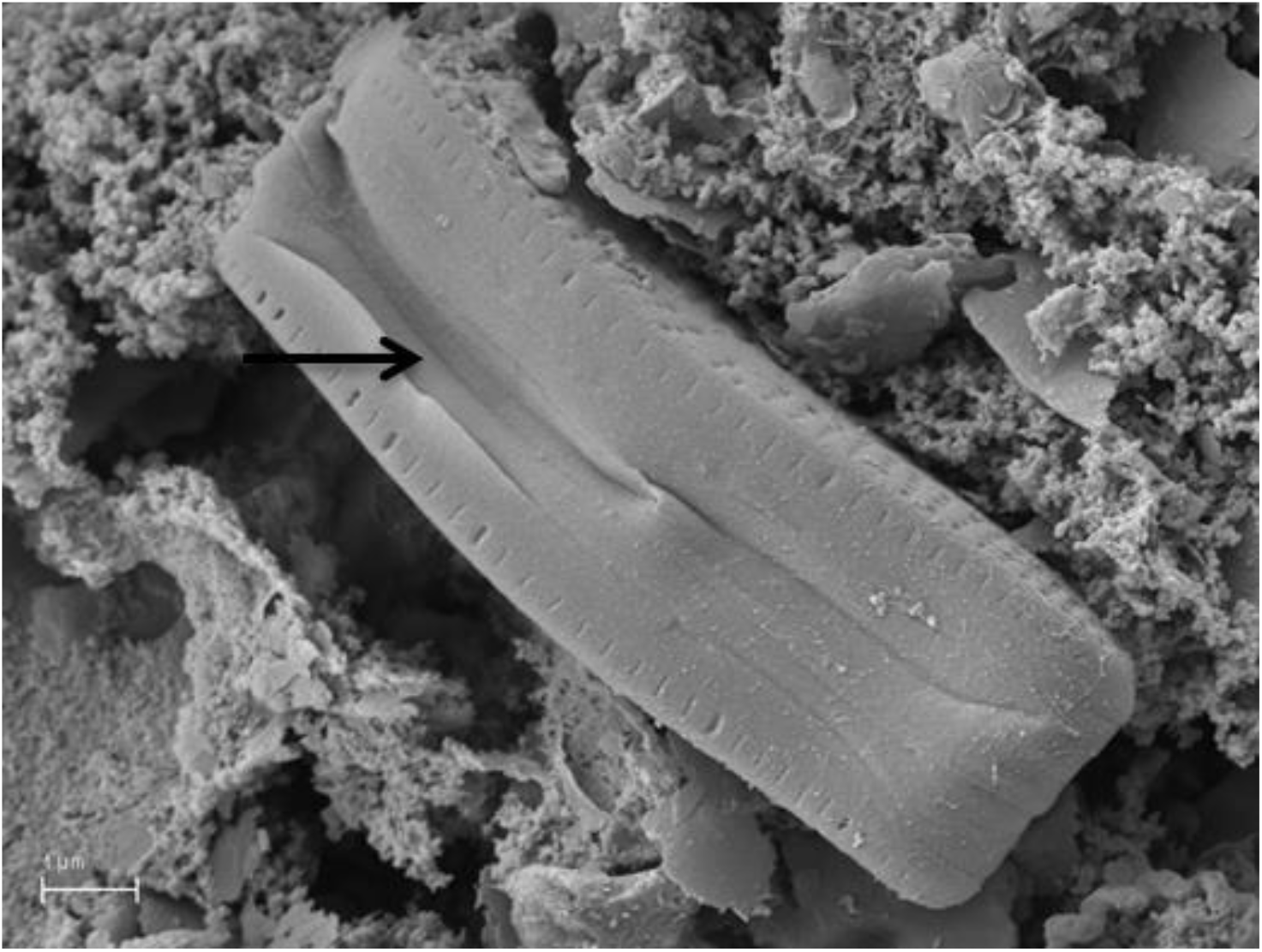
Deformed valvocopula seen at SEM (abnormal frustule identified in unprocessed sample).

Although the *Achnanthidium minutissimum* complex has been revised several times by many authors, insufficient information on its ecology is available. *Achnanthidium minutissimum* is omnipresent (Van Dam et al., 1994), often considered tolerant to severe water pollution (Stevenson and Bahls, 1999), but sometimes regarded as indicator of nutrient-poor waters (Potapova and Charles, 2007) or good water quality (Prygel and Coste, 1998). *Achnanthidium minutissimum* species complex are one of the most frequently occurring diatoms in freshwater and can be found in different environments, from unpolluted oligotrophic to highly polluted hypereutrophic waters (Hlúbiková et al., 2011). Many authors mention a positive correlation between *A. minutissimum* abundance and metal concentrations in contaminated sites (Cantonati et al., 2014). And Morin et al., 2008b led to the conclusion that *A. minutissimum* can be considered highly tolerant to metal contamination, producing teratologies only as a response to specific chemical contaminants (Falasco et al., 2009b).

Two alternative hypotheses can be proposed to explain this kind of teratology:

a. The girdle is poorer in silica and softer than the valve (Francius et al., 2008), so that it becomes flexible (Cox, 2011). The extent of silicification of a girdle band gradually diminishes from the region near the raphe to the overlap region (Kröger et al., 2007). Since metal contamination increases the rate of valve size diminution (characteristic of diatom asexual reproduction), valve surface does not decrease as rapidly as the cell volume (Falasco et al., 2009a). Frustule growth is only possible by parental valve separation synchronized with formation of new girdle bands (Stantos, 2010), so that the new girdle bands formed may not fit in the resulting frustules, adopting an aberrant outline. Moreover, diatoms exposed to metals might be increasing the number of girdle bands, due to the increase of vacuoles size, forcing chloroplast to the frustule edges (Gonçalves, 2017).
b. Teratology in diatoms has been explained in terms of malfunctions of proteins involved in silica transport and deposition (Falasco, 2009a). A family of these proteins, called cingulins, are specifically located in the girdle-band region of the frustule, where they are confined to the proximal surface of the terminal girdle bands of one of the valves (De Sanctis et al., 2016), and are presumed to be involved in the adhesion between girdle bands (Kröger and Wetherbee, 2000) and girdle band synthesis (Shrestha et al., 2012). It has been suggested that silicic acid uptake by diatoms via cingulins is mediated by a Zn-dependent system (Jaccard et al., 2009). A Zn excess affects the biochemical pathway of silicon metabolism (Martin-Jézéquel et al., 2003) and, particularly, metal-induced alteration of cingulins may impair girdle functioning (Karp-Boss et al., 2014).

We suggest a lab experimental approach in order to elucidate the nature of this teratology and interpret its occurrence in diatom populations in environmental terms.

## Acknowledgements

The present contribution was financially supported by a grant of the Romanian National Authority for Scientific Research, CCCDI – UEFISCDI, project 3-005 Tools for sustainable gold mining in EU (SUSMIN).

Thanks are also due to the research group from Resources and Ecosystems Area from Catalan Institute for Water Research (ICRA), Girona, Spain, for all their support and facilities.

## References

1. Cantonati M, Angeli N, Virtanen L, Wojtal AZ, Gabrieli J, Falasco E, Lavoie I, Morin S, Marchetto A, Claude F, Smirnova S (2014). *Achnanthidium minutissimum* (Bacillariophyta) valve deformities as indicators of metal enrichment in diverse widely-distributed freshwater habitats. Science of the Total Environment, 475: 201–215.

2. Cox EJ (2011). Morphology, cell wall, cytology, ultrastructure and morphogenetic studies. In The diatom world: 21–45. Springer Netherlands.

3. De Sanctis S, Wenzler M, Kröger N, Malloni WN, Sumper M, Deutzmann R, Zadravec P, Brunner E, Kremmer W, Kalbitzer HR (2016). PSCD domains of Pleuralin-1 from the diatom *Cylindrotheca fusiformis*: NMR structures and interactions with other biosilica-associated proteins. Structure, 24(7): 1178–1191.

4. European Standard EN 14407 (2004). Water quality – Guidance standard for the identification, enumeration and interpretation of benthic diatom samples from running waters. European Committee for Standardization, Brussels.

5. Falasco E, Bona F, Badino G, Hoffmann L, Ector L (2009a). Diatom teratological forms and environmental alterations: a review. Hydrobiologia, 623: 1–35.

6. Falasco E, Bona F, Ginepro M, Hoffmann L, Ector L (2009b). Morphological abnormalities of diatom silica walls in relation to heavy metal contamination and artificial growth conditions. Water sa, 35(5): 595–606.

7. Francius G, Tesson B, Dague E, Martin–Jézéquel V, Dufrêne YF (2008). Nanostructure and nanomechanics of live *Phaeodactylum tricornutum* morphotypes. Environmental microbiology, 10(5): 1344–1356.

8. Gonçalves SIB (2017). Zinc and copper impacts on freshwater diatoms: physiological, biochemical and metabolomic response of *Tabellaria flocculosa*. Master’s thesis. Universidade de Aveiro.

9. Hlúbiková D, Ector L, Hoffmann L (2011). Examination of the type material of some diatom species related to *Achnanthidium minutissimum* (Kütz.) Czarn. (Bacillariophyceae) Algological Studies, Volume 136/137: 19–43.

10. Hofmann G, Werum M, Lange-Bertalot H (2011). Diatomeen im Süβwasser-Benthos von Mitteleuropa. Koeltz Scientific Books, P.O. Box 1360, D-61453 Königstein, Germany. Hydrobiologia, 623: 1–35.

11. Jaccard T, Ariztegui D, Wilkinson KJ (2009). Incorporation of zinc into the frustule of the freshwater diatom Stephanodiscus hantzschii. Chemical Geology, 265(3): 381–386.

12. Johnson LM, Rosowski JR (1992). Valve and band morphology of some freshwater diatoms. V. Variations in the cingulum of *Pleurosira laevis* (Bacillariophyceae). Journal of Phycology, 28(2): 247–259.

13. Karp-Boss L, Gueta R, Rousso I (2014). Judging diatoms by their cover: variability in local elasticity of *Lithodesmium undulatum* undergoing cell division. PloS one, 9(10).

14. Kroger N, Poulsen N (2007). Biochemistry and Molecular Genetics of Silica Biomineralization in Diatoms. Handbook of Biomineralization: Biological Aspects and Structure Formation: 43–58.

15. Kröger N, Wetherbee R (2000). Pleuralins are involved in theca differentiation in the diatom *Cylindrotheca fusiformis*. Protist, 151(3): 263–273.

16. Lavoie I, Lavoie M, Fortin C (2012). A mine of information: benthic algal communities as biomonitors of metal pollution leaching from abandoned tailings. Sci Total Environ, 425: 231–241.

17. Lavoie I, Hamilton PB, Morin S, Tiam SK, Kahlert M, Gonçalves S, Falasco E, Fortin C, Gontero B, Heudre D, Kojadinovic-Sirinelli M, Manoylov K, Pandey LK, Taylor JC (2017). Diatom teratologies as biomarkers of contamination: Are all deformities ecologically meaningful?. Ecological Indicators, 82: 539–550.

18. Lechner CC, Becker CF (2015). Silaffins in silica biomineralization and biomimetic silica precipitation. Marine drugs, 13(8): 5297–5333.

19. Martin-Jézéquel V, Lopez PJ (2003). Silicon–a central metabolite for diatom growth and morphogenesis. Prog Mol Subcell Biol, 33: 99–124.

20. Morin S, Coste M, Hamilton PB (2008a). Scanning electron microscopy observations of deformities in small pennate diatoms exposed to high cadmium concentrations J Phycol, 44: 1512–1518.

21. Morin S, Duong TT, Boutry S, Coste M (2008b). Modulation de la toxicité des métaux vis-à-vis du développement des biofilms de cours d’eau (bassin versant de Decazeville, France)/Mitigation of metal toxicity to freshwater biofilms development (Decazeville watershed, SW France) Cryptogam Algol, 29 (3): 201–216.

22. Olenici A, Blanco S, Borrego-Ramos M, Momeu L, Baciu C (2017). Exploring the effects of acid mine drainage on diatom teratology using geometric morphometry. Ecotoxicology, 26(8): 1018–1030.

23. Ordinul Nr. 161/16.02.2006 al Ministerul Mediului și Gospodăririi Apelor pentru aprobarea Normativului privind clasificarea calității apelor de suprafață în vederea stabilirii stării ecologice a corpurilor de apă, Monitorul Oficial, Nr.511/2006.

24. Potapova M, Charles DF. 2007. Diatom metrics for monitoring eutrophication in rivers of the United States. Ecol. Indicators 7:48–70.

25. Prygiel J, Coste M (1998). Mise au point de l’indice Biologique Diatomée, un indice diatomique pratique applicable au réseau hydrographique français. L’Eau, l’Industrie, les Nuisances 211:40–5.

26. Santos JADC (2010). Cadmium effects in *Nitzschia palea* frustule proteins. Master’s thesis. Universidade de Aveiro.

27. Shrestha RP, Tesson B, Norden-Krichmar T, Federowicz S, Hildebrand M, Allen AE (2012). Whole transcriptome analysis of the silicon response of the diatom Thalassiosira pseudonana. BMC genomics, 13(1): 499 p.

28. Smol JP (2008). Pollution of lakes and rivers: a paleoenvironmental perspective (2nd ed.), Blackwell Publishing, Oxford: 383 p.

29. Stevenson RJ, Bahls LL (1999). Periphyton protocols. In M. T. Barbour, J. Gerritsen, B. D. Snyder and J. B. Stribling [Eds.] Rapid Bioassessment Protocols for Use in Streams and Wadeable Rivers: Periphyton, Benthic Macroinvertebrates and Fish. 2nd ed. EPA 841-B-99-002. US Environmental Protection Agency, Office of Water, Washington, DC, chap. 6.

30. Van Dam, H., A. Mertens and J. Sinkeldam (1994). A coded checklist and ecological indicator values of freshwater diatoms from the Netherlands. Neth. J. Aquat. Ecol. 28:117–133.

